# Module analysis using single-patient differential expression signatures improve the power of association study for Alzheimer’s disease

**DOI:** 10.1101/2020.01.05.894931

**Authors:** Jialan Huang, Dong Lu, Guofeng Meng

## Abstract

The causal mechanism of Alzheimer’s disease is extremely complex. It usually requires a huge number of samples to achieve a good statistical power in association studies. In this work, we illustrated a different strategy to identify AD risk genes by clustering AD patients into modules based on their single-patient differential expression signatures. Evaluation suggested that our method could enrich AD patients with common clinical manifestations. Applying it to a cohort of only 310 AD patients, we identified 175 AD risk loci at a strict threshold of empirical *p* < 0.05 while only two loci were identified using all the AD patients. As an evaluation, we collected 23 AD risk genes reported in a recent large-scale meta-analysis and found that 18 of them were re-discovered by association studies using clustered AD patients, while only three of them were re-discovered using all AD patients. Functional annotation suggested that AD associated genetic variants mainly disturbed neuronal/synaptic function. Our results suggested module analysis, even randomly clustering, helped to enrich AD patients affected by the common risk variants.

## 1 Introduction

Alzheimer’s disease is the most prevalent neurological disease among ageing population. It has been intensively studied for decades while its causal mechanisms remain elusive. Studies to the familial early-onset cases revealed a close association with three mutated genes, including APP, PSEN1 and PSEN2 [1]. They provided valuable insights into the contribution of amyloidogenic pathway as a causal mechanism of AD. Genome-wide association studies (GWAS) to late-onset AD patients discovered more rare and common risk variants. Among them, APOE *ε*4, an apolipoprotein, is the strongest genetic risk allele for late-onset AD, accounting for 3- (heterozygous) to 15-fold (homozygous) increase in AD risk [2]. However, it is still no clear how APOE contributes to AD genesis [3]. Many other risk genes, as listed in the AlzGene database (http://www.alzgene.org), are also discovered by GWAS. This entangles more biological processes and pathways as the risk mechanism of AD, such as immune system process (TNF, IL8, CR1, CLU, CCR2, PICALM and CHRNB2), cellular membrane organization (SORL1, APOE, PICALM, BIN1 and LDLR) and endocytosis (PICALM, BIN1, CD2AP) [4]. However, identified AD risk genes only explain a limited proportion of heritability, which indicates the complexity of AD genesis. Such diverse functional involvements of AD risk genes complicate mechanism studies. It is still a great challenge on how to illustrate the AD causal mechanism in an integrated way, limiting their application in new drug discovery.

Power is a critical consideration in association studies to detect risk variants [5]. As an extremely complex disease, AD often requires a large sample size to achieve a good power [3, 6]. For example, a recent meta-analysis included 71,880 cases and 383,378 controls, which identified 25 risk loci, implicating 215 potential causative genes [7]. However, such studies are limited by sample collection and cost, which blockades the discovery of more variants. To overcome such a problem, a strategy is to stratify patients based on some disease-relevant features [8]. For AD, carrier’s status of APOE-*ε*4 has been used to cluster AD population in association studies and reveals novel features [9]. Other factors, e.g. sex [10] and age [11], have also be used and the improved performance supports the values of population stratification in association studies.

Recently, many efforts were made to generate multi-omics data of AD for integrated studies. One example is the Accelerating Medicines Partnership - Alzheimer’s Disease (AMP-AD) projects, which includes transcriptomics, epigenomics, genetics, and proteomics data from over 2000 human brain samples. Some system biology analyses have been proposed for systematic insight into AD [12, 13, 14]. These studies led to a systematic understanding of how gene regulatory network perturbation contributed to the complex causal mechanism of AD and proposed key genes. However, such studies are also limited by the complexity of AD patients. The commonly used tools, such as WGCNA [15], MEGENA [16] and SpeakEasy [17], have limited consideration to population diversity. For complex diseases, e.g. AD, it is always a risk to treat the diverse patients as a homogeneous whole to compare with healthy controls. With the accumulation of multi-omics data, it allows a systematic integration of multiple omics data, e.g. to integrate genetic and transcriptomic data.

In this work, we proposed a new strategy to stratify AD patients based on the expression profiles similarity of single-patient DEGs. Our evaluation suggested that this method could enrich AD patients with common clinical manifestations. We applied it to 310 AD patients for both patient clustering analysis and genetic association studies. We identified 175 AD risk loci in 143 modules at a strict cutoff of empirical *p* < 0.05, while there were only two risk loci identified using all the AD patients. Function annotation suggested that identified risk genes were mainly related to neuronal/synaptic functions. We also evaluated 23 known AD risk genes and re-discovered 18 of them in at least one module. Allele frequency studies indicated that clustering analysis using single-patient DEGs enriched AD patients affected by common risk variants.

## 2 Results

### 2.1 A new pipeline to cluster AD patients utilizing single-patient DEGs

Considering the diversity of AD patients, we propose a new analysis strategy to cluster the AD patients affected by the common mechanisms. This method is based on differential expression analysis at single-patient levels. Figure 1(a) and Figure S1 describes the schema of the whole analysis pipeline. In our analysis, the reference expression profile was firstly built using the RNA-seq counts data of the normal individuals, which defined the ranges of gene expression values at a non-disease status. Next, gene expression values of patients were transformed into binary status by fitting to the reference expression profiles. In detail, if the gene expression values of patients exceeded the range of reference expression profiles, 1 or -1 is assigned to indicate up- or down-regulation. To improve confidence, a bi-clustering analysis algorithm is applied to perform filtering and cross-validation so that the whole set of single-patient differentially expressed genes (spDEGs) can be repeatedly observed in multiple patients, e.g. *n* = 5. Finally, using each patient as seed, we cluster the patients into modules if they carry the same set of spDEGs.

**Figure 1:**
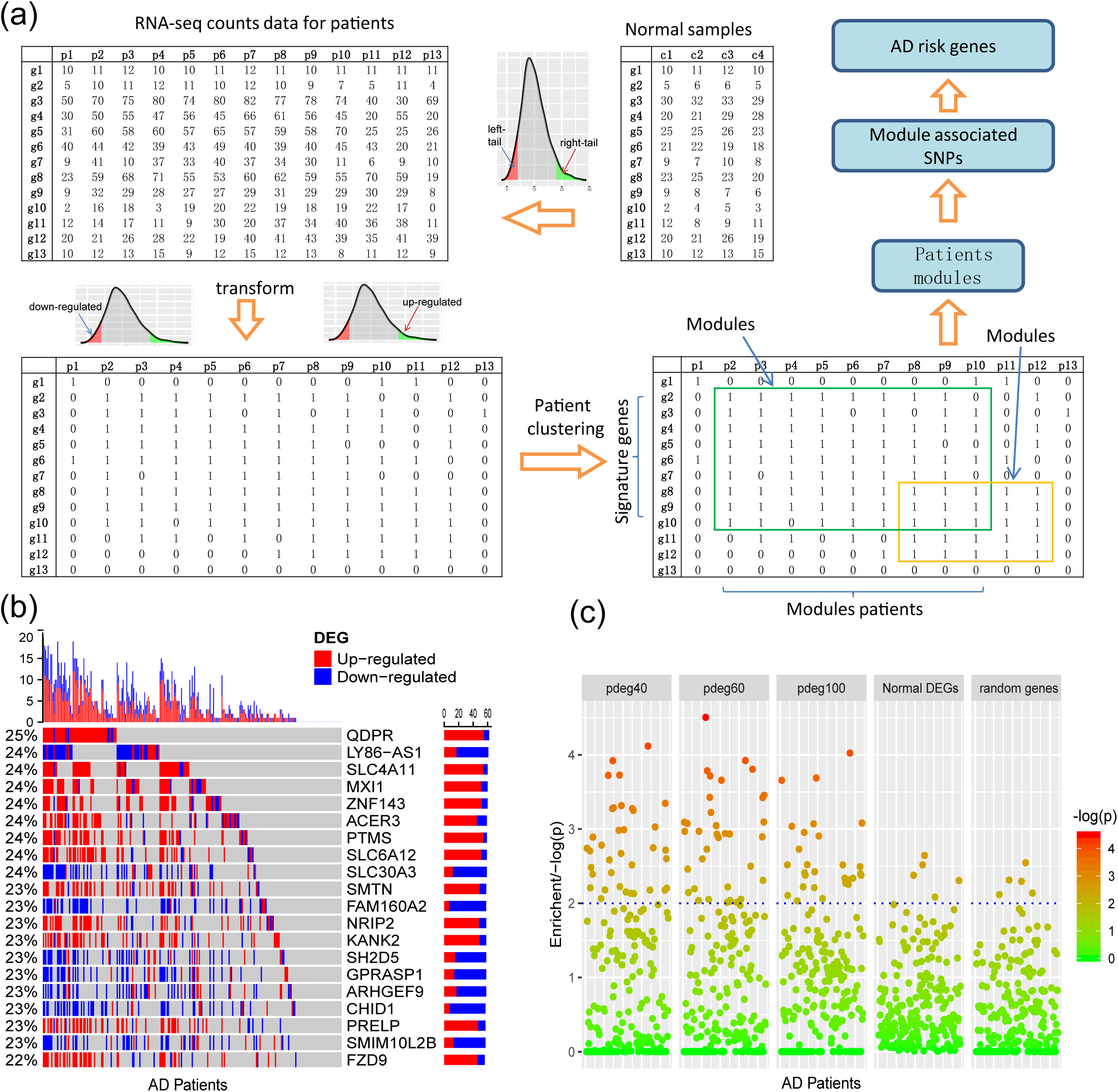
Clustering AD patients into modules based on single-patient differential expression profile similarity. (a) A analysis pipeline to cluster AD patients. The RNA-seq count data of AD patients were transformed into binary DEG matrix based on the reference profile built using the data of normal individuals; the AD patients with the shared DEG signatures are clustered as modules using a bi-clustering algorithm; genome-wide association study was performed in each patient module to identify the AD risk loci and genes. (b) Single-patient differential expression analysis indicated the complexity of AD patients, where genes displayed diverse DE status. (c) Clustering analysis enriched AD patients with similar clinical outcomes, e.g. cognitive test scores while not by the differentially expressed genes in all AD patients or random genes.

As an evaluation, we applied this pipeline to the dataset collected from the ROS/MAP study [2], which includes 251 AD samples with both RNA-seq data and clinical annotation. We identified cross-validated spDEGs for 171 patients. Among 15582 brain expressed genes, 3878 were predicted to be differentially expressed in at least one AD patient. Then, we investigated their differential expression status among all the AD patients. Figure 1(b) showed results of the top 20 most observed DEGs. We did not observe any shared differential expressed genes across all the AD patients. On the contrary, all DEGs were only differentially expressed in a small proportion of 251 AD patients. Additionally, we also observed inconsistent differential expression directions. Taking QDPR gene an as example, it was up-regulated in 22% of AD patients while also down-regulated in 3% AD patients. The similar results were observed with other spDEGs (see Figure 1(b)). We also performed clustering analysis using the most observed differential expressed genes and observed distinct differential expression patterns (see Figure S2). All these results suggested that AD patients were greatly diverse and that it would be a risk to treat AD patients as a homogeneous whole in any analysis.

Next, we investigated if AD patient clustering could enrich AD patients with common clinical manifestations. We generated patient modules based on sgDEG expression profile similarity. The modules were set to have different sizes, e.g. 40, 60, 100, which could be denoted as pdeg40, pdeg60 and pdeg100, respectively. The patients within the same module were supposed to be affected by the common mechanisms. As control, we also generated modules using randomly selecting genes and DEGs identified by traditional differential expression analysis. Figure 1(b) showed the evaluation results using cognitive scores (cts). At a cutoff of *p* < 0.01, 37 “pdeg60” modules were enriched with detrimental cts scores while only five modules identified by common DEGs or random genes were enriched. The most significant *p*-value was up to *p* = 2.51 × 10^−5^ in the “pdeg60” module. On the contrary, no module in “common DEG” and “random gene” exceeded the significance of *p* = 0.001. This result suggested that modules analysis using spDEG better enriched AD patients with common clinical manifestations.

### 2.2 More risk variants were identified in AD patient modules

We collected genotyping data from “hbtrc” study [18], including 310 LOAD patients and 153 non-demented healthy controls. We performed genome-wide association study (GWAS) using all the AD patients. In this process, we performed permutation procedure for 1000 times to estimate empirical *p* values. We found only two loci to have significant association with AD at a cutoff of empirical *p* < 0.05. The significant SNPs included rs2405283 (*p* = 1.15 × 10^−7^) and rs769450 (*p* = 1.65 × 10^−6^) (see Figure 2(a)). rs769450 was mapped to the second intron of APOE gene, consistent with published reports about the critical roles of APOE.

**Figure 2:**
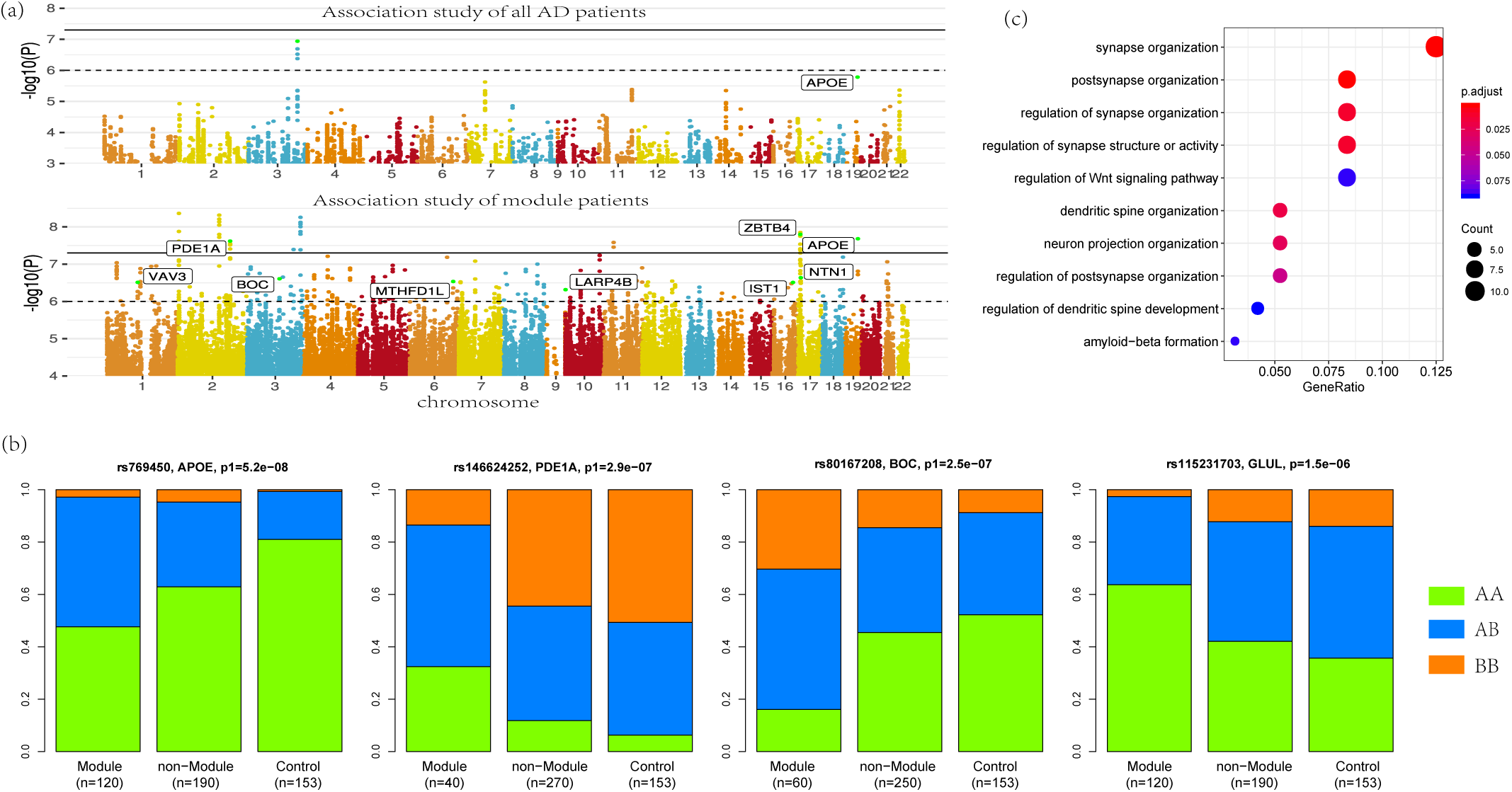
More risk variants were identified in AD patient modules. (a) Manhattan plot for the association studies to both all AD patients and patient modules, where more AD risk SNPs were identified in AD modules. (b) Allele frequencies in module patients, non-modules patients and control subjects. More risk allele enrichment was observed in module patients, suggested that clustering analysis enriched the AD patients affected by common risk variance. (c) Functional annotation to AD risk genes. Here, synaptic function related terms were most significantly enriched.

Applying clustering analysis, we predicted 143 modules of AD patients. Three association tests were performed for each module: (1) module patients against normal control; (2) module patients against non-module patients; and (3) non-module patients against normal control. The *p*-values were denoted as *p*1, *p*2 and *p*3, respectively. At a strict cutoff of empirical *p*1 < 0.05, we found 175 loci to have significant association in at least one of 143 modules (see Figure 2(a) and Table S1). Compared to association study using all the AD patients, more AD risk loci were observed within module patients. The APOE SNP rs769450 was observed in 41 modules and its association significance was also greatly improved. For example, the significance of rs769450 was up to *p*1 = 2.08 × 10^−8^ in a module of 80 AD patients while the significance for all 310 patients was *p*1 = 1.65 × 10^−6^. Tests between module patients and non-module patients supported allele frequency differences in 165 out of 175 loci at a cutoff of *p*2 < 0.01. Figure 2(b) showed the allele frequency for some exemplary SNPs. We observed that allele frequencies of identified risk SNPs were obviously different from the non-module patients and normal individuals. In most cases, non-module patients usually had similar allele frequencies with normal subjects. We checked if module patients were more associated with risk SNPs than non-module patients by comparing *p*1 and *p*3 value distribution (see Figure S3). We found module patients tended to report more significant association than non-module patients. It suggested that clustering analysis enriched the AD patient affected by the common risk SNPs.

We mapped 175 AD risk loci to 107 genes based on genomic proxy and GTEx eQTL annotation (see Table S1). Among them, 86 genes were observed in more than one module at a cutoff of empirical *p*1 < 0.05. APOE is the most observed risk gene, which is significantly associated with AD patients in 41 modules. We searched the published GWAS results and found that 46 genes had been reported for AD or brain-related function (see Table 1). Some of them had been reported in association studies of AD, such as PDE1A, JAM3, DLGAP1, CYYR1, SERPINB11 and MCPH1. To understand their function involvement, we performed Gene Ontology enrichment analysis to 107 AD risk genes (see Figure 2(c)). We found that the most enriched terms were also related to synaptic and neuronal function, e.g. “synapse organization” (*p* = 7.65 × 10^−6^). It suggested that the identified AD risk genes were related to normal brain function and had potential roles in AD genesis.

**Table 1:**
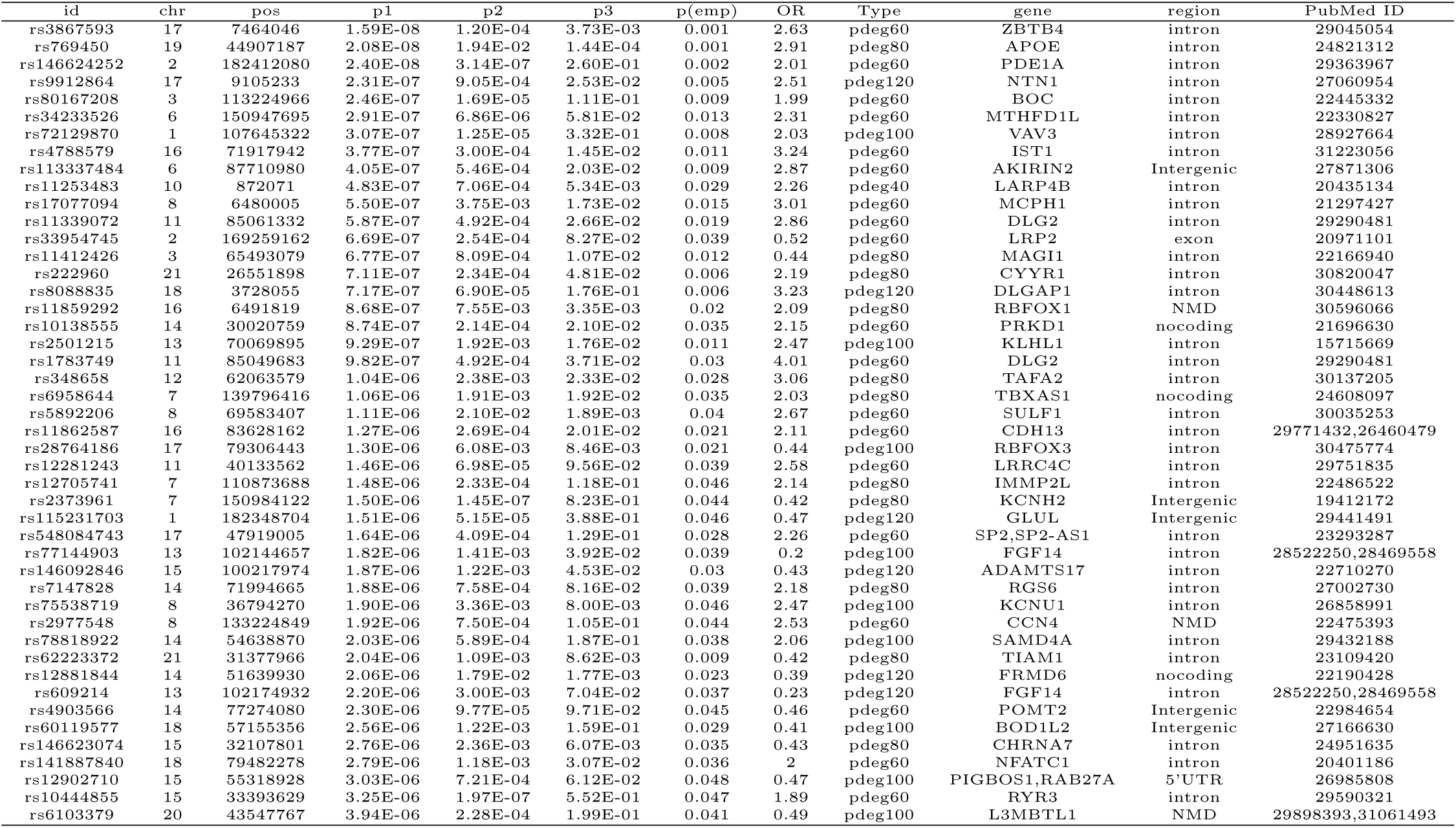
The results of association studies using module patients, non-module patients and control

In a recent large-scale meta-analysis, 23 AD risk loci were reported [6]. We checked their association using either all patients or module patients. We loosed the cutoffs of significant association by replacing empirical *p* < 0.05 with *p*1 < 10^−4^. Association study using all AD patients failed to identify any extra known AD risk gene to satisfy threshold of *p*1 < 10^−4^. Dislike the results using all AD patients, we observed that 18 out of 23 AD genes to have significant association with AD in at least one module. Table 2 summarized analysis results using module patients. By checking *p*2 and *p*3 values, we found significant allele frequency differences between module patients and no-module patients, supporting a conclusion that clustering analysis enriched AD patients affected by common known risk variants.

**Table 2:** The association results for known AD risk genes

| Association of module patients |  |  |  |  |  |  |  |  | Association of all AD patients |  |  |  |
| --- | --- | --- | --- | --- | --- | --- | --- | --- | --- | --- | --- | --- |
| SNP | gene | p1 | p2 | p3 | p(emp) | OR | module type | region | SNP | p1 | p(emp) | region |
| rs769450 | APOE | 2.08E-08 | 1.94E-02 | 1.44E-04 | 0.001 | 3.68 | pdeg80 | intron | rs769450 | 1.65E-06 | 0.015 | intron |
| rs71454394 | MS4A2 | 9.25E-06 | 3.73E-03 | 4.32E-02 | 0.257 | 2.48 | pdeg40 | intergenic | - | - | - | - |
| rs9462659 | TREM2 | 1.08E-05 | 8.99E-03 | 4.85E-02 | 0.35 | 2.02 | pdeg40 | intergenic | - | - | - | - |
| rs7152488 | SLC24A4 | 1.21E-05 | 1.85E-04 | 1.71E-01 | 0.175 | 0.3 | pdeg100 | intron | - | - | - | - |
| rs5021727 | HLA-DRB1 | 1.59E-05 | 1.80E-04 | 3.88E-01 | 0.389 | 0.45 | pdeg120 | intergenic | - | - | - | - |
| rs144409358 | CR1 | 2.09E-05 | 1.44E-03 | 1.79E-01 | 0.552 | 0.3 | pdeg120 | intron | - | - | - | - |
| rs12416009 | ECHDC3 | 2.10E-05 | 2.66E-04 | 2.19E-01 | 0.514 | 1.86 | pdeg40 | intergenic | - | - | - | - |
| rs9897336 | ACE | 2.41E-05 | 2.03E-04 | 4.91E-01 | 0.306 | 0.48 | pdeg100 | intergenic | - | - | - | - |
| rs55662472 | EPHA1 | 2.61E-05 | 5.33E-03 | 7.65E-02 | 0.519 | 3.15 | pdeg80 | intergenic | - | - | - | - |
| rs34708229 | MEF2C | 2.81E-05 | 4.09E-03 | 2.17E-02 | 0.675 | 2.45 | pdeg40 | intron | rs79820174 | 1.40E-04 | 1 | intron |
| rs6099038 | CASS4 | 2.86E-05 | 1.55E-04 | 3.16E-01 | 0.305 | 2.30 | pdeg100 | intergenic | - | - | - | - |
| rs13422890 | BIN1 | 3.35E-05 | 4.42E-06 | 8.04E-01 | 0.753 | 1.96 | pdeg60 | intron | - | - | - | - |
| rs36057699 | PTK2B | 3.39E-05 | 8.08E-03 | 4.63E-02 | 0.576 | 0.41 | pdeg120 | intron | rs36057699 | 8.70E-04 | 1 | intron |
| rs659023 | PICALM | 6.53E-05 | 8.73E-06 | 4.35E-01 | 0.797 | 0.54 | pdeg120 | intergenic | - | - | - | - |
| rs77792633 | FERMT2 | 8.95E-05 | 5.18E-04 | 5.44E-01 | 0.8 | 0.62 | pdeg60 | intergenic | - | - | - | - |
| rs57816367 | CD2AP | 9.17E-05 | 9.36E-05 | 4.60E-01 | 0.957 | 2.13 | pdeg40 | intron | - | - | - | - |
| rs10539341 | INPP5D | 9.42E-05 | 7.99E-03 | 9.36E-02 | 0.983 | 0.42 | pdeg100 | intron | - | - | - | - |
| rs2285898 | ABCA7 | 9.09E-05 | 1.00E-02 | 1.48E-01 | 0.632 | 0.53 | pdeg120 | intergenic | - | - | - | - |

### 2.3 Biological relevance of AD risk genes

Module based clustering analysis allows us to bridge AD risk genes to clinical features and affected biological processes. The clinical association of modules is determined by statistical test between module and non-module patients. Using HBRTC’s dataset, we identified nine and eight modules to be associated with braak and brain generalized atrophy at a cutoff of *p* < 0.01, respectively. Among them, 3 modules were associated with both braak and brain atrophy. Association study to these modules identified 8 and 20 loci respectively. In Table 3, we summarized the analysis results. These results supported that some AD risk genes might be more associated with some AD clinical outcomes. For example, NTN1 gene is a microtubule-associated force-producing protein and it is predicted to related to braak stage.

**Table 3:** The association results for known AD risk genes

SNPs associated with atrophy
| SNP | chr | pos | p1 | p1(emp) | seed patient | tp | gene | region | braak | atrophy |
| --- | --- | --- | --- | --- | --- | --- | --- | --- | --- | --- |
| rs147216627 | 1 | 157467609 | 7.04E-07 | 0.022 | X15888 | pdeg100 | - |  | 7.4E-02 | 9.05E-04 |
| rs78818922 | 14 | 54638870 | 2.03E-06 | 0.038 | X15888 | pdeg100 | SAMD4A | intron | 7.4E-02 | 9.05E-04 |
| rs1231702 | 11 | 29525814 | 4.97E-07 | 0.012 | X15914 | pdeg60 | AC110058.1,AC090124.1 | Intergenic | 0.95 | 7.96E-03 |
| rs3867263 | 18 | 63664376 | 9.40E-07 | 0.031 | X15914 | pdeg40 | SERPINB11 | intron | 0.24 | 7.18E-03 |
| rs236111 | 20 | 5952889 | 7.13E-07 | 0.011 | X15914 | pdeg60 | MCM8 | intron | 0.95 | 7.96E-03 |
| rs7113161 | 11 | 16969038 | 9.68E-07 | 0.024 | X15941 | pdeg120 | PLEKHA7 | intron | 1.34E-02 | 3.82E-03 |
| rs10489293 | 1 | 172217647 | 1.12E-07 | 0.005 | X16020 | pdeg40 | DNM3 | intron | 6.52E-02 | 7.87E-04 |
| rs12819631 | 12 | 104013393 | 2.84E-07 | 0.01 | X16020 | pdeg40 | GLT8D2 | intron | 6.52E-02 | 7.87E-04 |
| rs9912864 | 17 | 9105233 | 2.89E-06 | 0.037 | X16020 | pdeg100 | NTN1 | intron | 0.97 | 2.28E-03 |
| rs6875561 | 5 | 121537532 | 1.47E-06 | 0.049 | X16037 | pdeg80 | - |  | 1.27E-02 | 1.42E-03 |
| rs7930638 | 11 | 5567722 | 1.85E-06 | 0.043 | X16179 | pdeg120 | AC104389.4 | NMD | 3.56E-02 | 4.16E-03 |
| rs548084743 | 17 | 47919005 | 9.62E-07 | 0.021 | X16179 | pdeg40 | SP2,SP2-AS1 | intron | 9.51E-02 | 7.26E-03 |
| rs764624 | 14 | 71993857 | 2.32E-06 | 0.049 | X16183 | pdeg60 | RGS6 | intron | 0.11 | 8.66E-03 |
| rs78641850 | 10 | 53421383 | 2.17E-07 | 0.001 | X21821 | pdeg100 | - |  | 2.02E-02 | 6.43E-03 |
| rs17112518 | 14 | 21948703 | 2.30E-06 | 0.027 | X21901 | pdeg120 | - |  | 0.10 | 6.98E-03 |
| rs12881844 | 14 | 51639930 | 2.06E-06 | 0.023 | X21901 | pdeg120 | FRMD6 | Intergenic | 0.10 | 6.98E-03 |
| rs12480378 | 20 | 3110711 | 2.29E-06 | 0.025 | X21901 | pdeg120 | UBOX5-AS1,UBOX5 | Intergenic | 0.10 | 6.98E-03 |

SNPs associated with braak
| SNP | chr | pos | p1 | p1(emp) | seed patient | tp | gene | region | braak | atrophy |
| --- | --- | --- | --- | --- | --- | --- | --- | --- | --- | --- |
| rs6103379 | 20 | 43547767 | 3.94E-06 | 0.041 | X15917 | pdeg100 | Z98752.3,L3MBTL1 | NMD | 8.37E-03 | 1.01E-01 |
| rs11850894 | 14 | 22312243 | 2.04E-06 | 0.033 | X15989 | pdeg80 | TRAV40 | Intergenic | 1.33E-04 | 7.33E-02 |
| rs73699762 | 7 | 57341624 | 1.01E-06 | 0.028 | X15989 | pdeg120 | - |  | 2.55E-03 | 6.59E-02 |
| rs222960 | 21 | 26551898 | 4.39E-06 | 0.033 | X16038 | pdeg60 | CYYR1,CYYR1-AS1 | intron | 7.20E-03 | 5.03E-01 |
| rs6880404 | 5 | 163990493 | 9.32E-07 | 0.031 | X16105 | pdeg120 | - |  | 2.86E-03 | 4.77E-02 |
| rs538060878 | 17 | 9142309 | 1.10E-07 | 0.004 | X21810 | pdeg40 | NTN1 | intron | 4.27E-03 | 6.56E-01 |
| rs1016268 | 12 | 129517265 | 1.88E-06 | 0.048 | X21810 | pdeg80 | TMEM132D | intron | 6.58E-04 | 1.92E-01 |
| rs6769967 | 3 | 44217312 | 1.77E-07 | 0.012 | X21810 | pdeg40 | - |  | 4.27E-03 | 6.56E-01 |
| rs16885931 | 6 | 22265940 | 8.04E-07 | 0.021 | X21810 | pdeg120 | CASC15 | Intergenic | 6.11E-04 | 2.32E-02 |

SNPs associated with both atrophy and braak
| SNP | chr | pos | p1 | p1(emp) | seed patient | tp | gene | region | braak | atrophy |
| --- | --- | --- | --- | --- | --- | --- | --- | --- | --- | --- |
| rs769450 | 19 | 44907187 | 7.12E-07 | 0.008 | X16149 | pdeg120 | APOE | intron | 9.62E-03 | 1.94E-03 |
| rs78415808 | 12 | 69406115 | 8.07E-07 | 0.023 | X16183 | pdeg80 | - |  | 2.18E-03 | 5.84E-03 |
| rs820562 | 3 | 112745366 | 1.46E-06 | 0.042 | X16037 | pdeg120 | LINC02042 | Intergenic | 6.11E-03 | 3.99E-05 |

AD patient modules are always associated with a list of spDEG signature genes, which could be used to investigate biological relevance of AD risk genes. Figure 3 showed the analysis results of functional annotation to module spDEG signature genes. Among the significant terms, “extracellular matrix assembly”, “synaptic signaling”, “learning and memory” and “protein folding” were more observed or more significant. By textmining studies, we found many published evidence for their close association with AD, supporting that predicted AD risk genes contributed to AD development. For example, extracellular matrix was observed to have significant changes during the early stages of AD [19] and extracellular matrix could induce *β*-Amyloid Levels [20]. Among predicted risk genes, APOE, POMT2, FGF14, CDH13 and RBFOX3 display more functional involvements.

**Figure 3:**
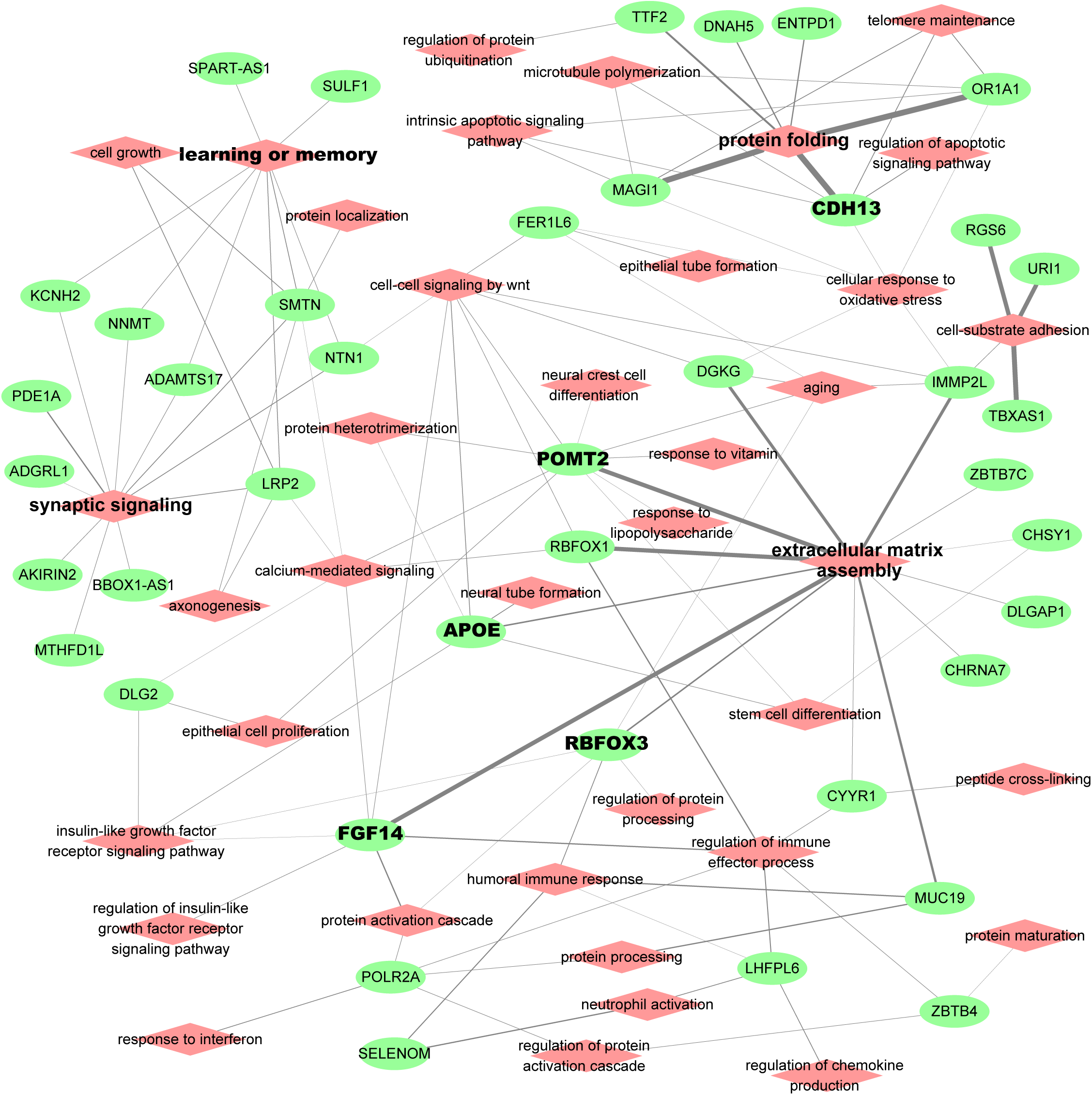
Functional relevance of AD risk genes. Here, the module spDEG signatures were used for Gene Ontology enrichment to indicate the functional involvement of modules.

### 2.4 Evaluation using randomly clustering of AD patients

In above analysis, we attempted to cluster AD patients with a common set of spDEGs so that the clustered patients were more affected by common AD variants. As an evaluation, we performed a simulated study by randomly splitting AD patients into simulated modules at corresponding sizes. Then we predicted AD risk SNPs using the exact same setting. In each round of simulation, we identified about 105 AD risk SNPs on average at a cutoff of empirical *p* < 0.05. We compared their analysis results to that of true modules and found that about 63% of risked SNP (out of total 175 loci) could be overlapped with the SNPs predicted using true modules. This evaluation seemed to support a conclusion that subsetting AD patients had benefits to improve the power of association studies, even when the criteria to stratify AD patients was to randomly pick up. Comparing to random modules, clustering using spDEG signature could recover more AD risk SNPs.

## 3 Discussion

In this work, we took more consideration to AD patient diversity and attempted to stratify patients into modules affected by different genetics background. Therefore, we came up with a analysis pipeline to cluster AD patients based on some assumptions, including that (1) AD patients are very diverse and differential expression patterns differ among AD patients; (2) we can used single-patient DEGs as biomarkers to indicate the dysregulation status of AD patients and to cluster the AD patients affected by common mechanisms. In our previous work, we have applied similar strategies to discover enriched transcription factor binding sites [21] and cancer driver mutations [22], and achieved good performance. Evaluation using real patient data suggested that this method could group AD patients with similar clinical outcomes and common risk variants, validating our assumptions.

We applied a new strategy to find the differentially expressed genes for each AD patient and clustering patients based on the spDEG signatures. In this process, we made some assumptions. For example, we defined the reference expression profiles for normal individuals by fitting to a Gaussian or negative binomial distribution. The robustness of this step was dependent on the number and homogeneity of control individuals. To identify the differentially expressed genes, we need to set some thresholds to determine if the gene expression level of one AD patient was beyond the normal ranges. In our work, we tested different cutoffs and selected *p* = 0.1.

We did association study in each module of size 40 to 120. Compared to the study using all AD patients, the statistical power decreased with decreased sample size in each association study. However, more AD risk loci were identified for increased number of AD patient modules. 175 loci were predicted to be associated with AD at a strict threshold of empirical *p* < 0.05, while only two loci exceed such a threshold using all AD patients. The genotype frequency was found to be different between module and non-module patients. All these results suggested that AD risk variants might contribute only limited subset of AD population.

In this work, we proved the benefits of patient clustering in association studies to AD. In our application, we reported more AD risk genes even when only 310 AD patients were used. In large-scale meta analysis, there were about 20-30 genes identified as AD risk genes [7, 23]. However, by searching public literature and databases, e.g. GWAS catalog, we found more than 100 studies and more than 300 genes that had been reported in associated studies to AD patients. These studies could be treated as a subset of large-scale AD meta-analysis. This results suggested that there might be more AD risk genes and AD patients subsetting helped to identify them.

## 4 Methods

### 4.1 The samples and subjects

The AD and control sample data were collected from the “ROS/MAP” study [2] and “HBTRC” study [18]. “ROS/MAP” data included the genotype, expression and clinical data for 1788 subjects. The AD-related clinical annotation were provided by the data suppliers. The important one included ages, the cognitive score (cts), years of education, ApoE genotype, braak stage (braak) and assessment of neuritic plaques (ceradsc). We use the clinical annotation for “cogdx”, a physician’s overall cognitive diagnostic category, to select the AD patient (cogdx = 4 or 5) and control subjects (cogdx = 1). After filtering the ones with missed or unclear information for either clinical records or RNA-seq, we found 219 AD patients and 187 control subjects that would be used for module analysis and clinical enrichment studies. “HBTRC” study had both RNA-seq and genotype data for 573 samples, including 311 AD samples. We filtered the one with missed clinical information, RNA-seq or genotyped data. Finally, 310 AD patients and 153 control subjects were used.

### 4.2 Clustering the AD patient using single-patient DEGs

We developed a computational algorithm to cluster AD patients (see Figure S1). The main idea behind this tool is that AD patients are highly diverse and can affected by divergent mechanisms; it is possible to cluster AD patients if they shared a subset of differentially expressed genes (DEGs). This algorithm is implemented in R package `DEComplexDisease`.

It mainly includes four steps:

- Utilize RNA-seq data of normal subjects to construct reference expression profiles. In this step, the parameters of negative binomial distribution or Gaussian distribution are estimated to describe the distribution profile of non-disease samples;
- The gene expression of AD patients are transformed into binary differential expression status. In this step, the expression values of genes are fitted against reference expression profiles. Binary differential expression status is assigned as 1, -1 or 0 to indicate up-, down-regulation or no difference;
- Apply a bi-clustering analysis to identify DEGs that are repeatedly observed in multiple AD patients, e.g. n=5;
- Using the spDEG of each AD patient as signature, we compute the co-expression correlation and identify the patients with the most similar expression profiles to construct modules.

The R codes are publicly available in https://github.com/menggf/DEComplexDisease.

### 4.3 Clinical manifestation association analysis

“ROS/MAP” data mainly includes three AD related clinical features, including cognitive score (cts), CERAD score and braaksc. “HBTRC” has clinical information for braak and atrophy. Such clinical features can be used to evaluate the disease relevance of modules. Therefore, we applied our tool to generate modules of different sizes, e.g. 40, 60 and 120. For each module, AD patients can be grouped as module patient and non-module patients. We did Kolmogorov-Smirnov (KS) test to evaluate the clinical manifestation differences between two groups of AD patients.

### 4.4 Processing genotype data

We applied stringent quality control (QC) filters to the genotype data. First, we removed the individuals with missing genotype rates > 0.05 and SNPs with missing call rate > 0.05. In next step, the SNPs with minor allele frequency MAF < 0.1 or Hardary-Weinberg equilibruim *p*-value < 1.0 × 10^−5^ were excluded. The individual with autosomal heterozygosity above empirically determined thresholds were filtered. Identity-by-descent (IBD) of all possible gene pairs were also calculated and we removed the ones with potential genetic relatedness. These QC filters were performed for multiple rounds to make sure that no indivivual or SNP could be filtered any way. Then, We performed prephasing in SHAPEIT2 [24] using the 1000 Genome Project data as reference. Then, we conducted whole-genome imputation using IMPUTE2 [25] in 5-Mb segments with a filtering of the SNP with MAF less than 0.1 in EUR population. The imputed data were evaluated for quality control using the thresholds mentioned before. We performed principal component analysis (PCA) on autosomal genotype data and adjustment for stratification.

### 4.5 Association study

Association studies were performed for both all AD patients and module patients. To simplify it, we only include the definite AD patients and control individuals in association analysis so that binary disease status could be assigned for each patient. We performed population stratification by use of the principal components of chromosomal genetic variations. Association analysis perform using fast score test implemented in `GenABEL` package. In this step, the first 10 principle components were used as covarites to remove the effects of population structure to make sure of no clear evidence of inflation in the association results. To control the false positive discovery, we also estimated the empirical *p*-values using performing permutation analysis by generating the distribution under the null hypothesis for 1000 times. In each round of call, minimal p-value was compared with original p values. For a SNP, its empirical *p*-values is defined as a proportion of times minimal p-values in 1000 resampling less than the original p-value. We set empirical *p*-values < 0.05 as the cutoff to select the module associated SNPs. The codes for association studies is available in https://github.com/menggf/spDEG_and_Association.

