## Supplementary figures and images for "Module analysis using single-patient differential expression signatures improve the power of association study for Alzheimer’s disease"

### Figure S1

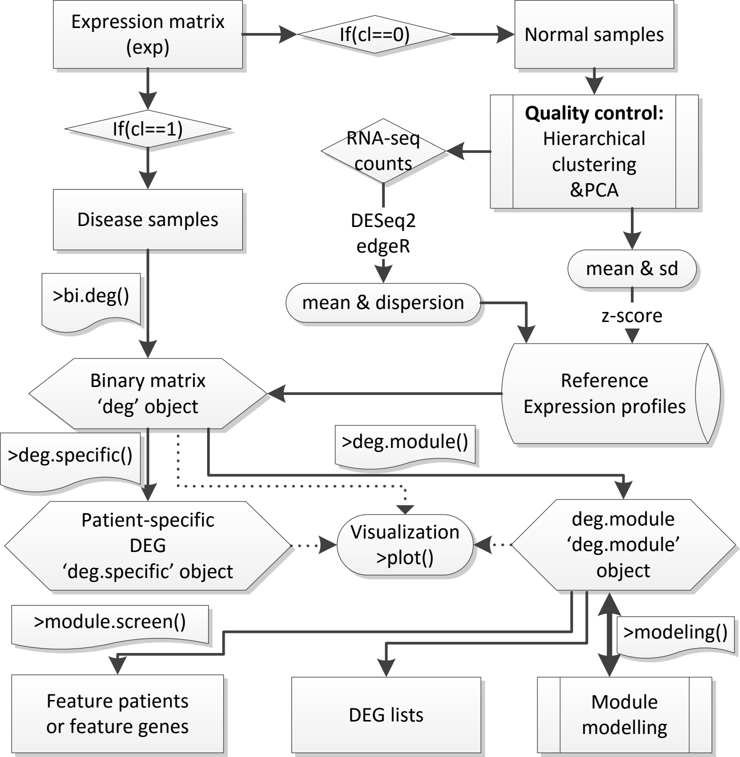

### Figure S2

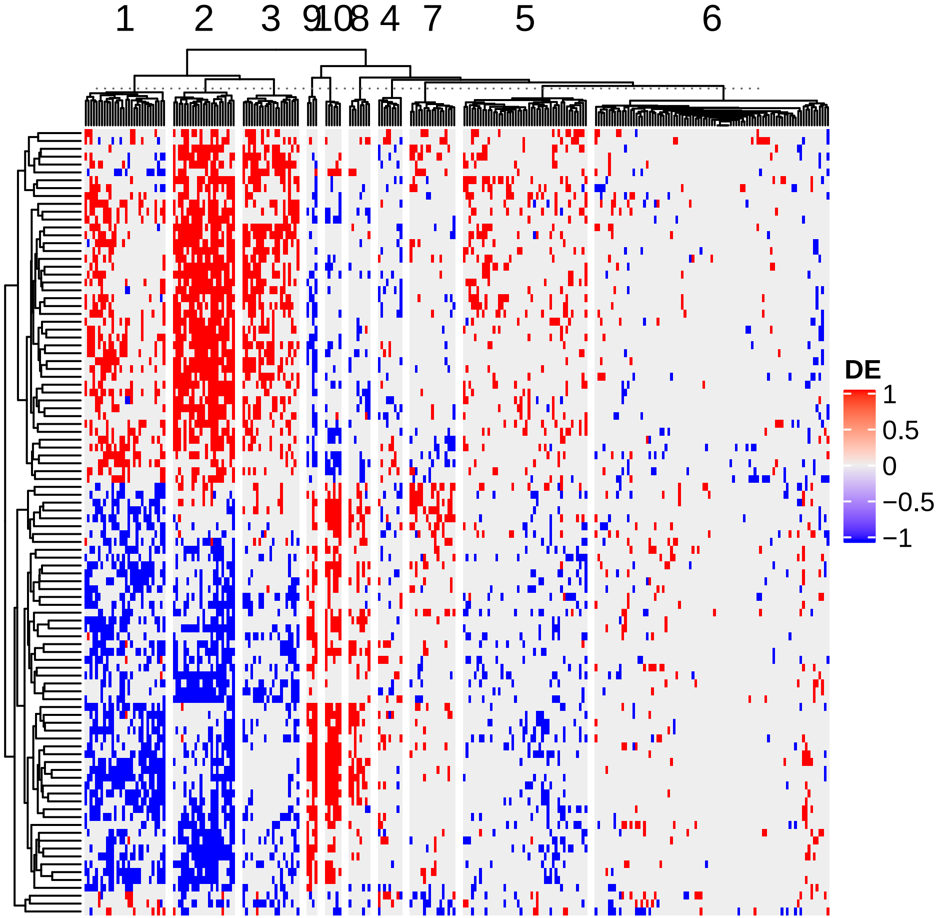

### Figure S3

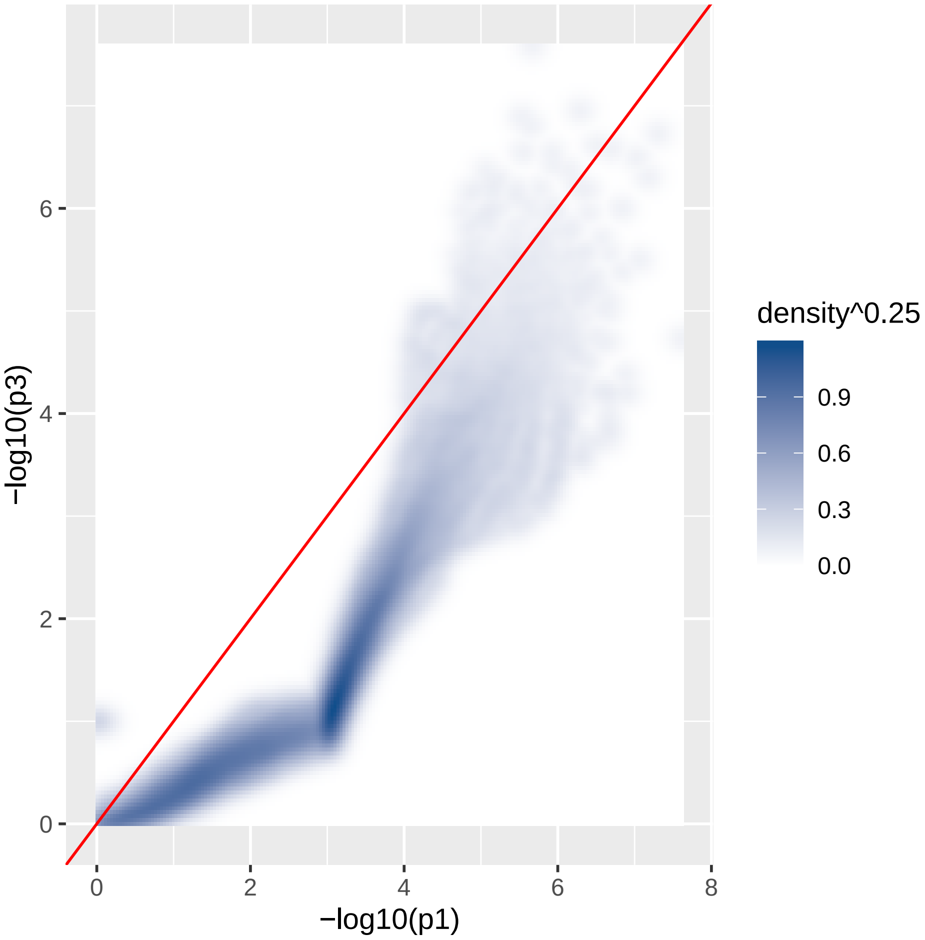
